# Decoding host-microbiome interactions through co-expression network analysis within the non-human primate intestine

**DOI:** 10.1101/2023.08.11.552617

**Authors:** Mika Uehara, Takashi Inoue, Sumitaka Hase, Erika Sasaki, Atsushi Toyoda, Yasubumi Sakakibara

**Author notes:** Corresponding Author Yasubumi Sakakibara, 3-14-1 Hiyoshi, Kohoku-ku, Yokohama, 223-8522, Japan.

## Abstract

The gut microbiome affects the health status of the host through complex interactions with the host’s intestinal wall. These host-microbiome interactions may spatially vary along the physical and chemical environment of the intestine, but these changes remain unknown. This study investigated these intricate relationships through a gene co-expression network analysis based on dual transcriptome profiling of different intestinal sites—cecum, transverse colon, and rectum— of the primate common marmoset. We proposed a gene module extraction algorithm based on graph theory to find tightly interacting gene modules of the host and the microbiome from a vast co-expression network. The 27 gene modules identified by this method, which include both host and microbiome genes, not only produced results consistent with previous studies regarding the host-microbiome relationships, but also provided new insights into microbiome genes acting as potential mediators in host-microbiome interplays. Specifically, we discovered associations between the host gene *FBP1*, a cancer marker, and polysaccharide degradation-related genes (*pfkA* and *fucI*) coded by *Bacteroides vulgatus*, as well as relationships between host B-cell specific genes (*CD19*, *CD22*, *CD79B*, *PTPN6*) and a tryptophan synthesis gene (*trpB*) coded by *Parabacteroides distasonis*. Furthermore, our proposed module extraction algorithm surpassed existing approaches by successfully defining more functionally related gene modules, providing insights for understanding the complex relationship between the host and the microbiome.

**IMPORTANCE:** We unveiled the intricate dynamics of the host-microbiome interactions along the colon by identifying closely interacting gene modules from a vast gene co-expression network, constructed based on simultaneous profiling of both host and microbiome transcriptomes. Our proposed gene module extraction algorithm, designed to interpret inter-species interactions, enabled the identification of functionally related gene modules encompassing both host and microbiome genes, which was challenging with conventional modularity maximization algorithms. Through these identified gene modules, we discerned previously unrecognized bacterial genes that potentially mediate in known relationships between host genes and specific bacterial species. Our findings underscore the spatial variations in host-microbiome interactions along the colon, rather than displaying a uniform pattern throughout the colon.

## INTRODUCTION

The mutual interaction between the intestinal microbiome and the host constitutes a complex biological process, with intertwined dependencies that are essential for the survival and health of both. The interactions between the intestinal microbiome and the host are formed by various factors such as the effects of metabolites produced by bacteria on the host, the binding of bacterial-derived proteins and host receptors, and the binding of bacterial cell surface structures and host receptors. Studies on host-microbiome interaction through bacterial-derived proteins have discovered that a novel protein P9 secreted by *Akkermansia muciniphila* binds to host ICAM-2, thereby increasing the secretion of glucagon-like peptide 1 (GLP-1), suggesting that the interaction between P9 and ICAM-2 could be a potential therapeutic target in type 2 diabetes patients (1). Excluding certain gram-negative bacteria, it has also been reported that bacterial lipopolysaccharides (LPS), which are components of the gram-negative bacterial cell wall, bind to the Toll-like receptor 4 (TLR4) complex and can induce innate immune responses (2). Among these interactions, the host-microbiome interaction mediated by metabolites is particularly well-studied. For instance, primary bile acids, produced in the host’s liver cells, are converted into secondary bile acids by the microbiome in the colon. This microbiome-derived secondary bile acid is known to regulate the homeostasis of regulatory T cells, which are immune-suppressive cells in the host (3). Through these processes, the intestinal microbiome and host mutually interact and establish a symbiotic relationship while forming a metabolic pathway or physiological response.

To understand such complex interactions between the host and the microbiome, it is beneficial to analyze combined profiles of both the microbiome and the host. Recent studies have reported on dual profiling of host and microbiome gene expression levels to understand the interactions between the host and microbiome (4, 5, 6). Previously, the focus was primarily on identifying bacterial species related to host phenotypes (such as diseases) based on the relative abundance of microbiome (7). However, profiling both host and microbiome gene expression levels has provided further insights into which genes of which bacterial species are related to host physiological functions and phenotypes. Gene co-expression network analysis is one of the useful approaches for comprehensively deciphering complex interactions from the vast number of gene expression profiles of the host and microbiome. A gene co-expression network is constructed by representing genes as nodes and connecting genes with correlated expression levels via edges. Representing gene relationships as a network enables the extraction of closely interconnected gene modules (subgraphs) and the identification of hub genes connected to multiple genes, thereby enhancing interpretability. For instance, a study (5) characterizing the host-microbiome interaction during influenza virus infection identified modules from the host’s gene co-expression network. It discovered correlations between host gene modules involved in interferon signal transduction and the sulfonamide antibiotic resistance genes of the microbiome. In another study (6), network analysis was applied to the gene expression profiles of the host and microbiome in the nasopharyngeal airway during bronchiolitis. Networks were constructed for both the host and microbiome and modules were identified from these networks. The study revealed that host modules were involved in regulating host-T-cell activity. Additionally, microbiome modules involved in microbial-BCAA metabolism and oxidative stress response were associated with disease severity.

Despite such progress, several limitations remain in understanding the interactions between the host and the intestinal microbiome. First, there’s a lack of understanding of the spatial variation in host-microbiome interactions. The colon, rich in microbiota, can be subdivided into the cecum, ascending colon, transverse colon, descending colon, sigmoid colon, and rectum. Each section possesses distinct functions and differs in its physicochemical environment, including variations in nutrient content, pH, and luminal flow (8, 9, 10) and the microbiome could vary as well. Indeed, studies using animal models have reported that the composition and functional fluctuation of the microbiome varies due to environmental changes depending on intestinal location (biogeographical location) (11, 12, 13). However, these studies focused solely on the microbiome, and the coordinated spatial variation of both the microbiome and host gut cells along the intestinal tract has yet to be clarified.

Second, there is a lack of computational methods for extracting significant modules (subgraphs) from the host-microbiome gene co-expression network. Analyzing a massive gene co-expression network integrating the gene expression profiles of both the host and microbiome helps interpret the interactions between the genes of the host and the microbiome. Especially in a huge gene co-expression network, it is crucial to extract densely connected subgraphs (modules) to divide the network into understandable units; however, existing algorithms for extracting modules from the network are insufficient for accurately capturing the interaction between the host and microbiome because they were not designed with the purpose of understanding interactions between organisms (14, 15, 16, 17).

Lastly, there are concerns regarding the applicability of the results of studies using animal models to humans. While mouse models have been the basis of biological research for a long time due to ethical constraints, experiments in controlled environments, and considerations of reproducibility, there are several issues and limitations to their use. The physiology, anatomy, and pharmacology of mice differ from those of humans, and there are differences in the composition and function of the intestinal microbiome between mice and humans (13, 18). Particularly considering the complexity of host-microbiome interactions, it is challenging to precisely replicate the relationship between humans and their intestinal microbiome. In contrast, the common marmoset, a small New World primate, is considered a useful model in preclinical trials because it is genetically close to humans and has common physiological and anatomical characteristics (19). Moreover, the common marmoset is the only non-human primate in which a germ-free state has been successfully achieved (20), and it has a fecal microbiota similar to that of humans compared to other animal models, including mice (13), potentially broadening the scope of intestinal microbiome research.

To address these challenges, this study aimed to reveal the interactions between the host and the microbiome along multiple sites of the common marmoset colon—cecum, transverse colon, and rectum—by developing a module extraction method applicable to the gene co-expression network that represents the complex interactions between the host and the microbiome. Through simultaneous profiling of host and microbiome transcripts in the intestinal sites of the common marmoset, we constructed an integrated gene co-expression network, incorporating both host and microbiome genes. From this network, we further identified densely covariant gene modules (subgraphs) to interpret the interactions between genes. As conventional methods for extracting gene modules from the network were insufficient for interpreting the interactions between organisms examined in this study, we proposed a module extraction algorithm based on graph theory. Our proposed method successfully extracted more modules containing functionally related genes of both the host and microbiome compared to existing algorithms, underlining its suitability for predicting host-microbiome interactions. This method extracted 27 gene modules containing both host and microbiome, indicating that the gut microbiome is functionally closely involved in the variation of host gene expression along the colon.

## RESULTS

### Overview of the analysis pipeline integrating host and microbiome transcriptomes: Identification of closely interacting gene modules from gene co-expression networks

To investigate the host and microbiome interactions along the colon, we proposed an analysis method integrating the host transcriptome and the microbiome metatranscriptome (Fig. 1). First, we computed gene expression profiles from the host transcriptome and microbiome metatranscriptome obtained via dual RNA-seq and created three types of gene co-expression networks: host-host, host-microbiome, and microbiome-microbiome. By unifying these three networks, we constructed a host and microbiome gene co-expression network, with the genes of the host and microbiome serving as nodes, and covariant genes interconnected. Second, from the gene co-expression network, we identified gene modules containing closely interacting genes from both the host and microbiome. We aimed to interpret the interactions between host genes and microbiome genes within modules extracted from a gene co-expression network, based on the assumption that functionally similar genes are co-expressed (21). Conventional module extraction methods are not designed to understand interspecies interactions (14, 15, 16, 17) and do not distinguish between host gene and microbiome gene nodes in the network. Consequently, modules composed solely of host genes or exclusively of microbiome genes can be extracted, making it challenging to understand the interactions between the host and the microbiome when such single species modules are present. Therefore, we proposed a clique-finding-based module extraction method that takes into account the structure of three networks: host-host, host-microbiome, and microbiome-microbiome. This method begins by exploring cliques from the gene co-expression network. In graph theory, a “clique” refers to a subgraph within the network where all nodes are fully connected. For overlapping cliques, the method calculates the cliqueness in each of the original three networks and merges them if the cliqueness of all networks exceeds a certain threshold, thereby obtaining the final gene modules (refer to the MATERIALS AND METHODS section for details).

**Fig. 1.**
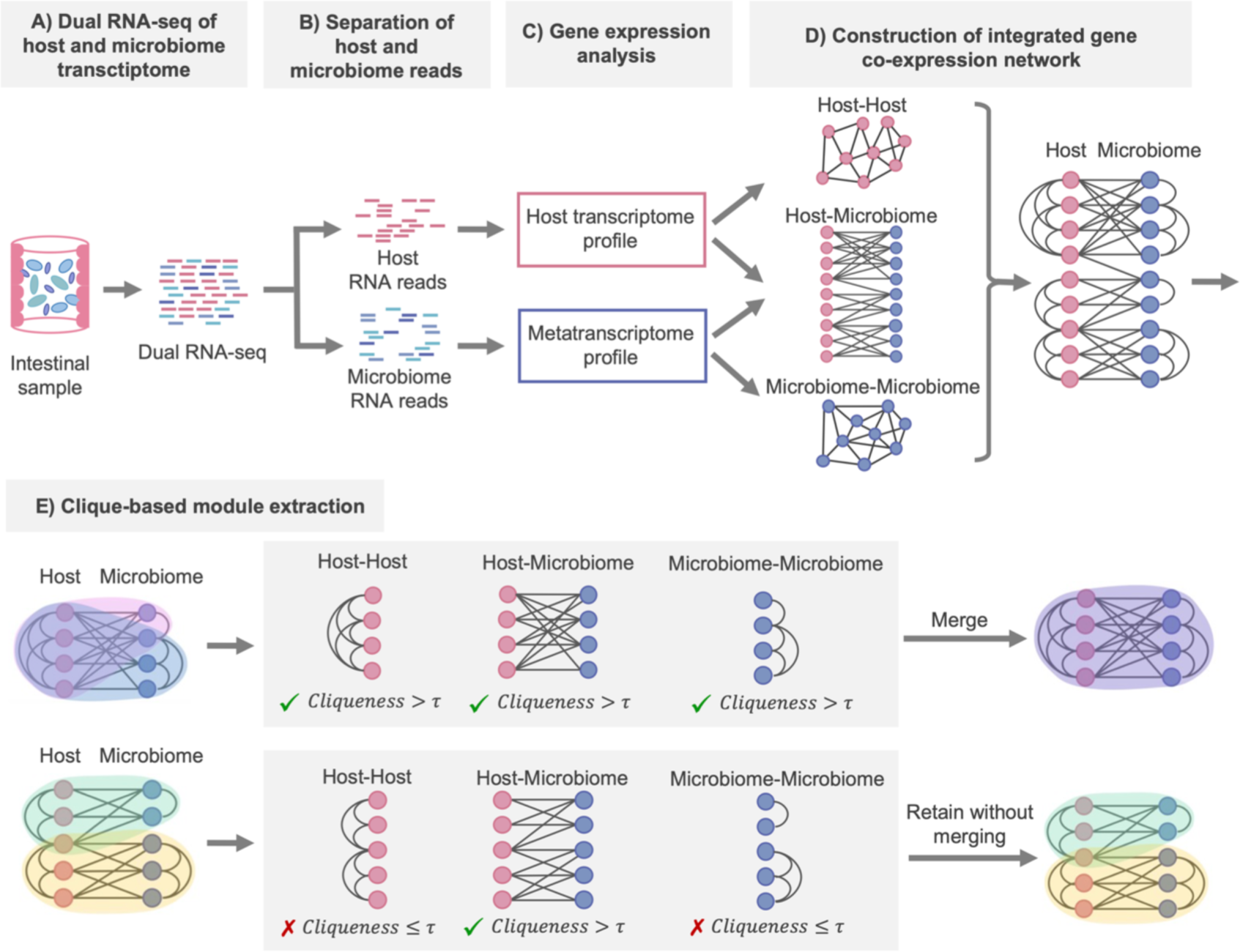
Overview of the main analytical approach used in this study. This method involves the simultaneous acquisition of the host and microbiome transcriptome (metatranscriptome) from intestinal sites and conducts a gene co-expression network analysis of the host and microbiome community. The procedure is outlined from step A to E. (A) Perform dual RNA-seq of host and microbiome transcriptomes in internal site samples. (B) Separate host and microbiome RNA reads in silico using the host reference genome. (C) Compute the gene expression profiles of the host and microbiome, respectively. (D) Unify gene co-expression networks between host-host, host-microbiome, and microbiome-microbiome. The host-host and microbiome-microbiome gene co-expression networks are connected when there is a positive correlation between genes, while the host-microbiome gene co-expression network is connected when there is either a positive or negative correlation between genes. (E) Identify closely interacting gene modules from the gene co-expression network using a proposed clique-based algorithm. (E Step 1) This process begins with searching for cliques. (E Step 2) If overlaps exist between cliques, the cliqueness score is computed in three gene co-expression networks (host-host, host-microbiome, and microbiome-microbiome) when overlapping cliques are merged. (E Step 3) If the cliqueness score exceeds the merging threshold τ in all three networks, the cliques are merged into a single module. If the score does not exceed the merging threshold τ in at least one network, the original cliques are retained without merging. This process is repeated until no further modules can be merged.

### Computing gene expression profiles of the host and the microbiome from dual RNA-seq data

First, we extracted high-quality RNA (RNA integrity number (RIN) scores ≥8.6; Table S1) from the intestinal sites (cecum, transverse colon, and rectum) of five common marmosets (Supplementary Note Section 1) and carried out dual RNA-seq with NovaSeq 6000 v1.5 (100 bp PE). The obtained RNA read data amounted to a total of 4,724 M reads (mean 157,450,033 paired-end reads/sample; Table S2; Supplementary Note Section 2).

Next, as the RNA read data obtained by dual RNA-seq contains a mixture of reads derived from both the host and the microbiome, it is necessary to separate these read data in silico (Table S3; Supplementary Note Section 3). The pipeline established in this study separates the host-derived read data and the microbiome-derived read data by aligning the RNA reads to the host reference genome (Fig. 1B). To avoid transcriptomic cross-contamination of read data between host and microbiome read data, we adjusted the pipeline parameters using simulated RNA read data from the host and the microbiome (Supplementary Note Section 4). As a result, we achieved a precision of 1.0 in separating the read data of the host and the microbiome, and our established pipeline completely separated the reads derived from the host and those from the microbiome (Table S9 posted at https://doi.org/10.5281/zenodo.10425960).

After the above preprocessing, we calculated the gene expression levels (transcripts per million (TPM)) for both the host and the microbiome. As a result, we detected 16,578 host expressed genes and 8,533 microbiome expressed genes in total (Supplementary Note Section 5 and 6). To examine the expression patterns across different sites and across different individuals for both host genes and microbiome genes, we applied Principal Component Analysis (PCA) to the gene expression profiles for visualization. There is a clear separation between the host gene expression profiles for each intestinal site, and the host samples were not grouped based on individuals (Figures 2A and B). The microbiome profiles at the cecum and rectum, which are the beginning and end of the colon, respectively, were separated by PC1, with samples from the transverse colon, situated between the cecum and rectum, spanning across both sites (Figure 2C). For the microbiome profiles as well as for the host ones, samples were not separated based on individuals (Figure 2D). These results suggest that both host and microbiome gene expression profiles are more variable across different sites compared to those across individuals.

**Fig. 2.**
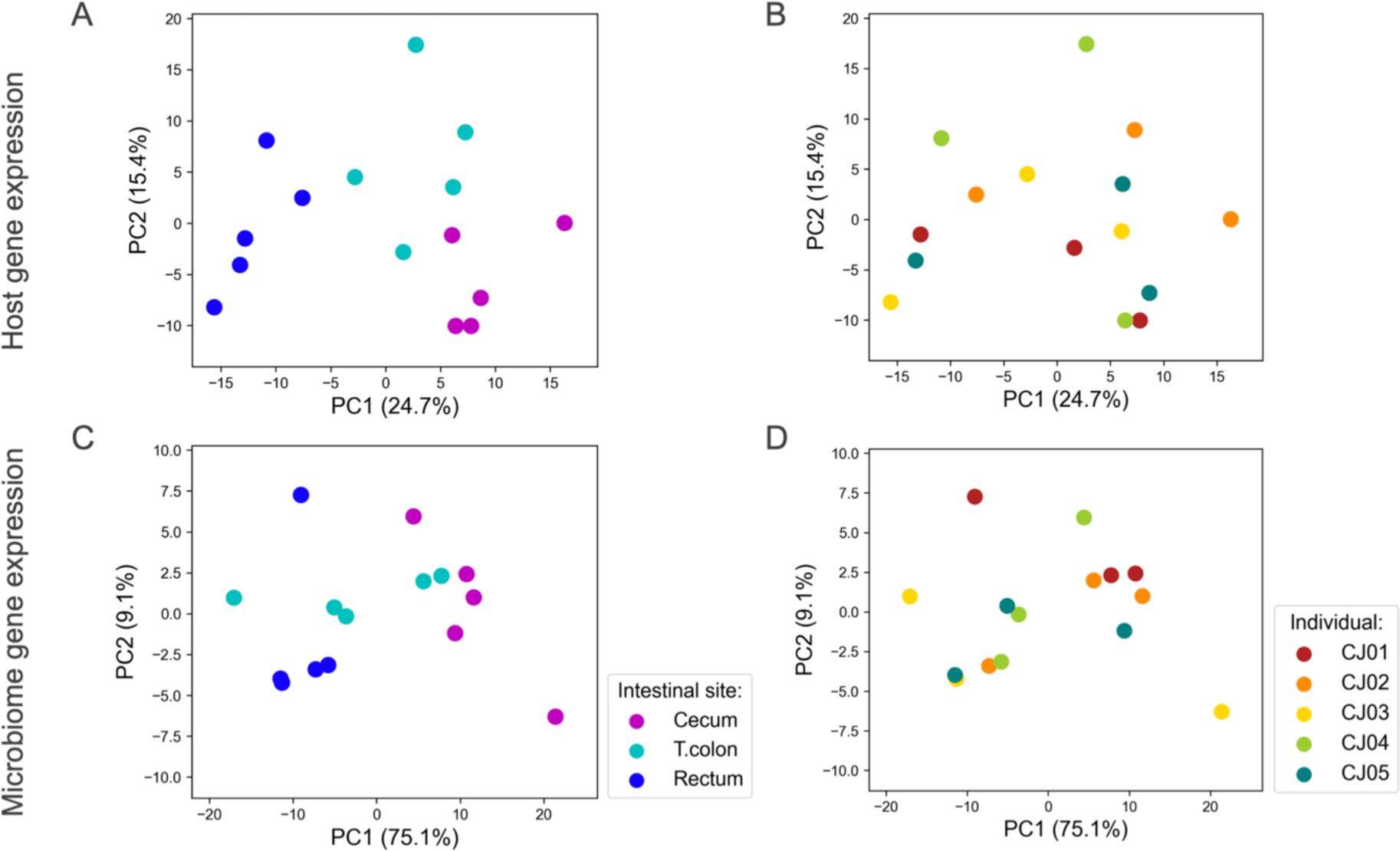
PCA analysis of gene expression data for host and microbiome. PCA was applied to the host gene expression profiles to visualize groupings according to site (A) and individual (B). PCA was similarly applied to the microbiome gene expression profiles to visualize groupings according to site (C) and individual (D).

### Spatial variations in host gene expression among intestinal sites

Of the 16,578 host genes detected in three regions of the common marmoset’s intestinal tract— cecum, transverse colon, and rectum—a total of 2,615 genes showed significant expression variations among these sites (FDR < 0.01) (Table S4; Supplementary Note Section 5).

Furthermore, upon conducting functional pathway analysis, 40 pathways were identified, all of which were significantly upregulated in the cecum compared to the transverse colon (normalized enrichment score > 1.5, FDR < 0.1; Table S10 posted at https://doi.org/10.5281/zenodo.10425960). In particular, pathways with high normalized enrichment scores, such as Fc epsilon RI signaling pathway, T cell receptor signaling pathway, leukocyte transendothelial migration, Yersinia infection, and B cell receptor signaling pathway were detected, indicating that pathways related to immune functions were upregulated in the cecum relative to the transverse colon (Fig. 3A). Additionally, a substantial number of genes involved in these functional pathways were found to be significantly upregulated in the cecum (FDR < 0.01). For example, among the genes specifically upregulated in the cecum, we found genes related to B cell receptor signaling. These include *CD19*, which acts as a coreceptor for the B cell antigen receptor complex and augments B cell activation (22); *CD79A* and *CD79B*, components of the B cell antigen receptor that generate signals upon antigen recognition (23); and *CD22* and *CD72*, which inhibit the signal transduction of the B cell antigen receptor (24, 25). In addition, genes associated with T cell receptor signaling were also found as being highly expressed specifically in the cecum. These include *CD28*, a co-stimulatory receptor for T cell receptors that activates T cells (26); *CD3D*, a component of CD3 that plays a role in signal transduction during T cell activation (27); *CD40LG*, which is induced on the T cell surface following T cell receptor (TCR) activation (28); and *CTLA4*, which attenuates T cell activation and fine-tunes immune responses (29) (Fig. 3B).

**Fig. 3.**
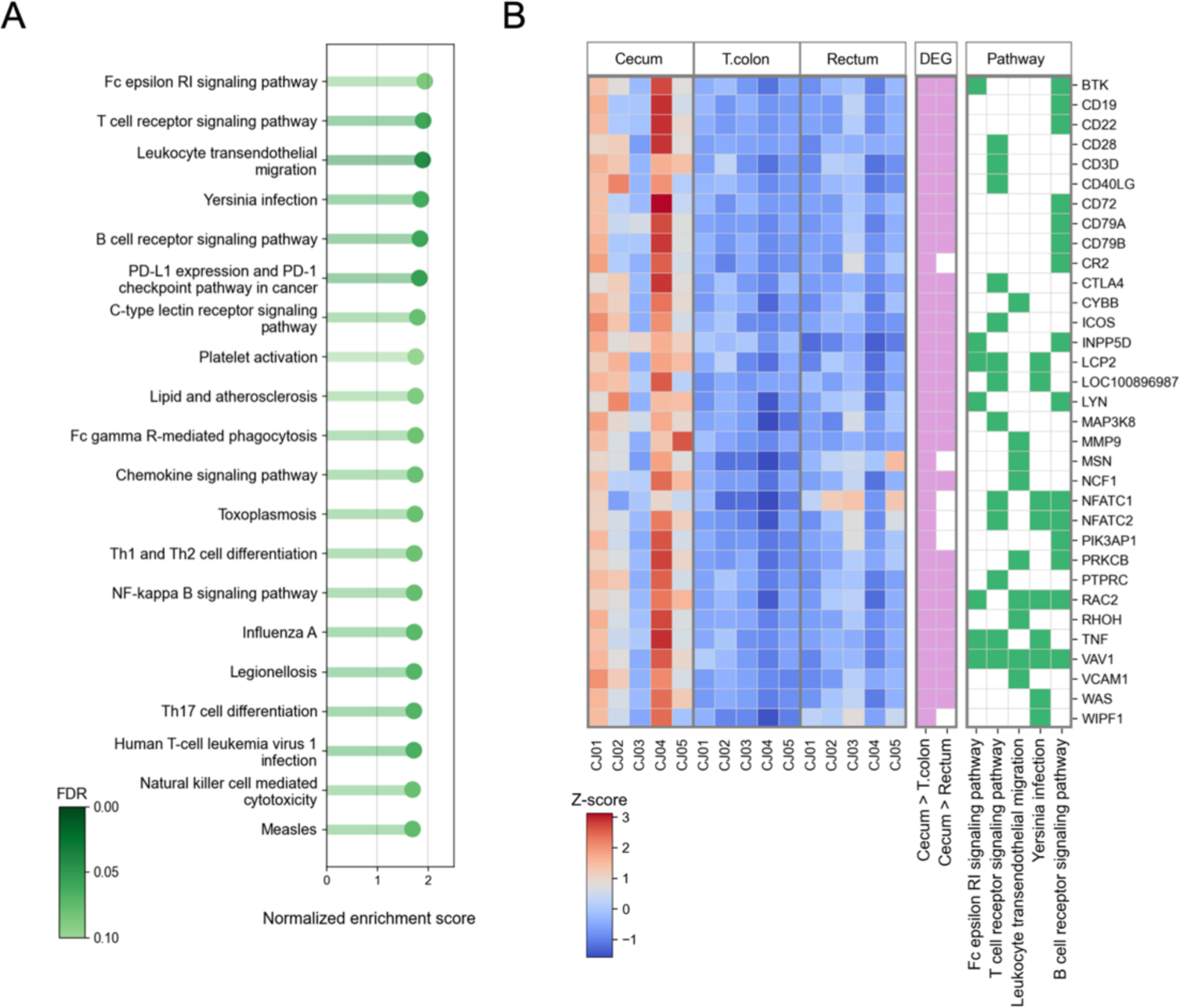
Variations in host gene expression and functional pathways among intestinal sites. (A) The top 20 functional pathways significantly upregulated in the cecum compared to the transverse colon (normalized enrichment score > 1.5, FDR < 0.1), as revealed by gene set enrichment analysis, are presented alongside their normalized enrichment score and FDR. (B) Heatmap of differentially expressed genes belonging to the top 5 enriched functional pathways. The heatmap on the left represents the z-scores of gene expression levels, with genes significantly upregulated in the cecum compared to the transverse colon and rectum (FDR < 0.01) colored in pink on the middle heatmap. The heatmap on the right indicates the functional pathways to which the genes belong, represented in green.

### Interpreting host-microbiome interactions through spatial gene co-expression network analysis within the intestine

We constructed the unified gene co-expression network composed of 1,359 gene nodes, which embodies co-expression information on host-host, host-microbiome, and microbiome-microbiome interactions (Table S11, S12, and S13 posted at https://doi.org/10.5281/zenodo.10425960; Supplementary Note Section 7). Subsequently, to identify densely co-expressed gene sets within this network, we implemented the clique-based module extraction algorithm proposed in this study. As a result, we then identified 27 gene modules containing both host and microbiome genes (Table S14 and S15 posted at https://doi.org/10.5281/zenodo.10425960). Furthermore, upon conducting a functional pathway enrichment analysis on these gene modules, we found that 70.4% (19 out of 27) of gene modules could be functionally characterized (Supplementary Note Section 8).

The modules we identified were characterized by functional pathways including the B cell receptor signaling pathway, fructose and mannose metabolism, oxidative phosphorylation, and biosynthesis of amino acids (Fig. 4). In addition, we provided visualizations of the bacterial species that contribute to microbiome gene expression within these identified modules, together with host and microbiome genes (Fig.5, 6, 7 and 8).

**Fig. 4.**
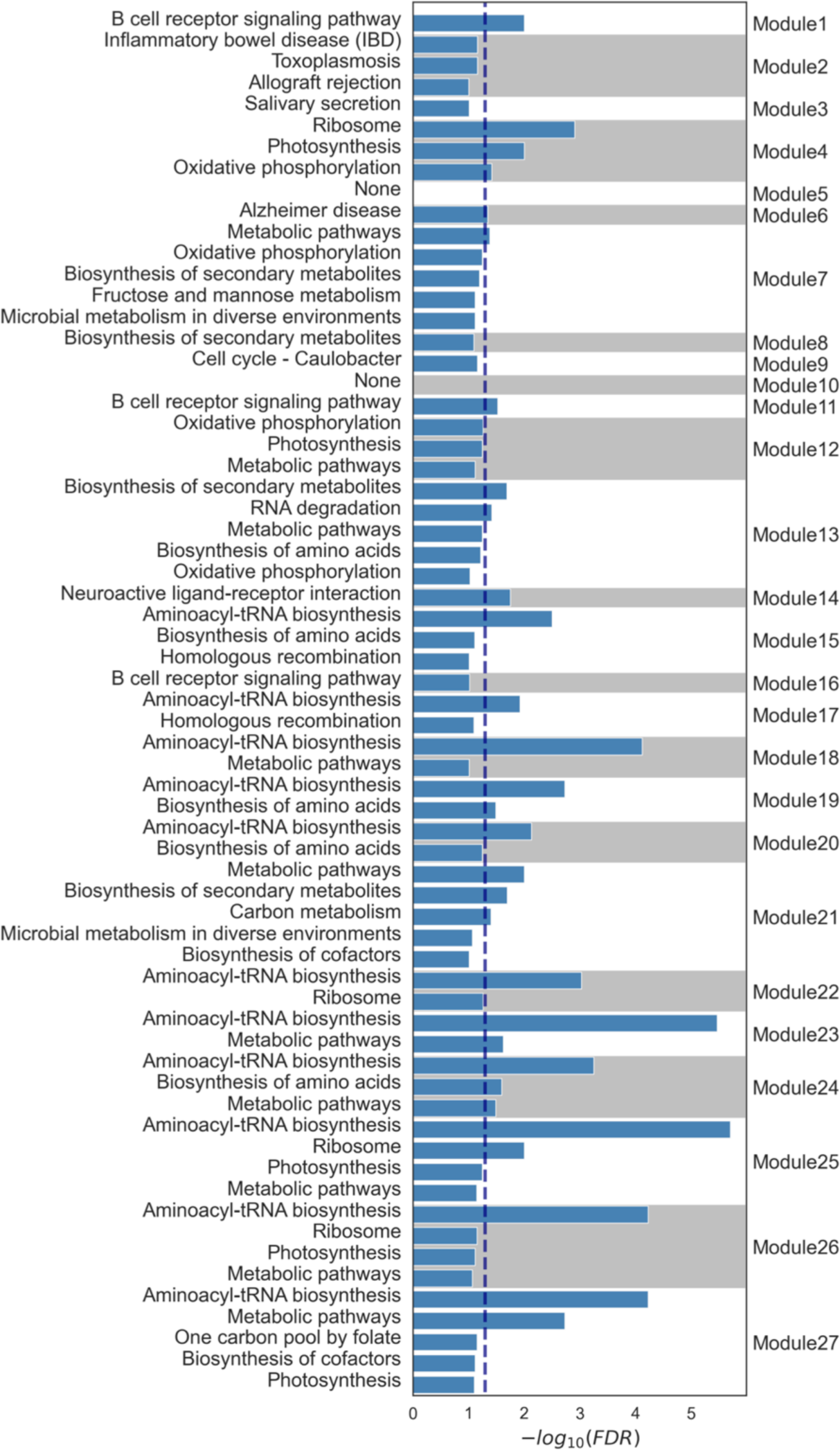
Functional pathways characterized in modules extracted from the host-microbiome gene co-expression network. This figure displays the identified modules along with the results of their functional pathway enrichment analysis (Supplementary Section 8). To enhance clarity, the background of the bar graphs alternates between gray and white for each module. Modules were characterized using KEGG pathways, with those meeting an FDR < 0.05 as determined by Fisher’s exact test. The dashed line represents an FDR < 0.05 (-log10(FDR) > 1.30). All pathways with an FDR < 0.1 are represented, and modules with no identified function are denoted as “None”.

**Fig. 5.**
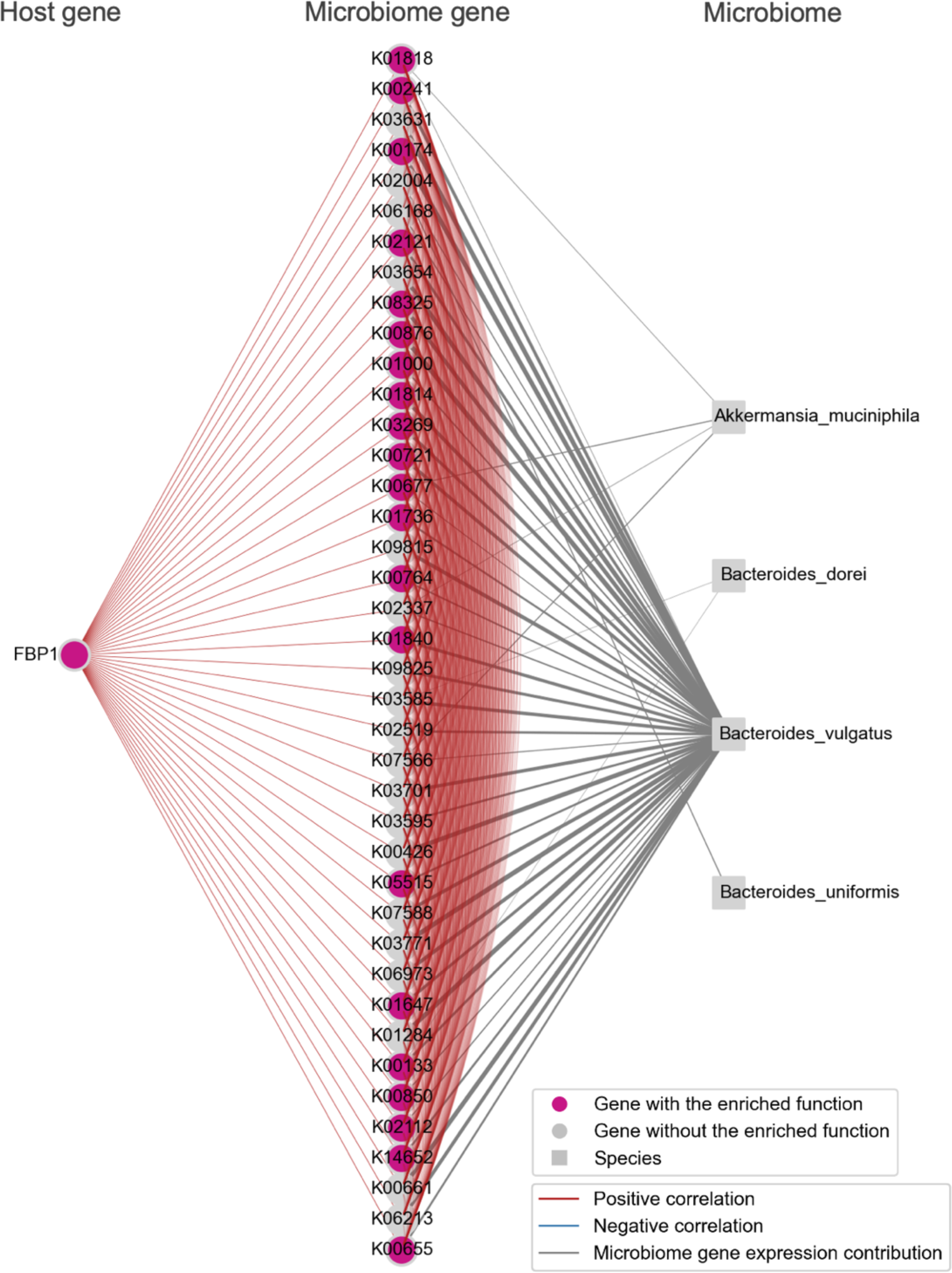
Metabolic pathway-enriched Module_07. A module showing a positive correlation between the host gene *FBP1*, which is highly expressed in the transverse colon, and 40 microbiome genes, including those involved in fructose-mannose metabolism and oxidative phosphorylation, was extracted. These genes were visualized alongside bacterial species nodes that contribute to the expression of the microbiome genes. Note that the bacterial species nodes were added after the module detection analysis. Edges among genes indicate an absolute Spearman’s correlation coefficient greater than 0.8. Edges connecting microbiome genes to bacterial species are shown when the contribution rate of bacterial species to microbiome gene expression exceeds 0.2. The thickness of the edge increases with higher contribution rates.

**Fig. 6.**
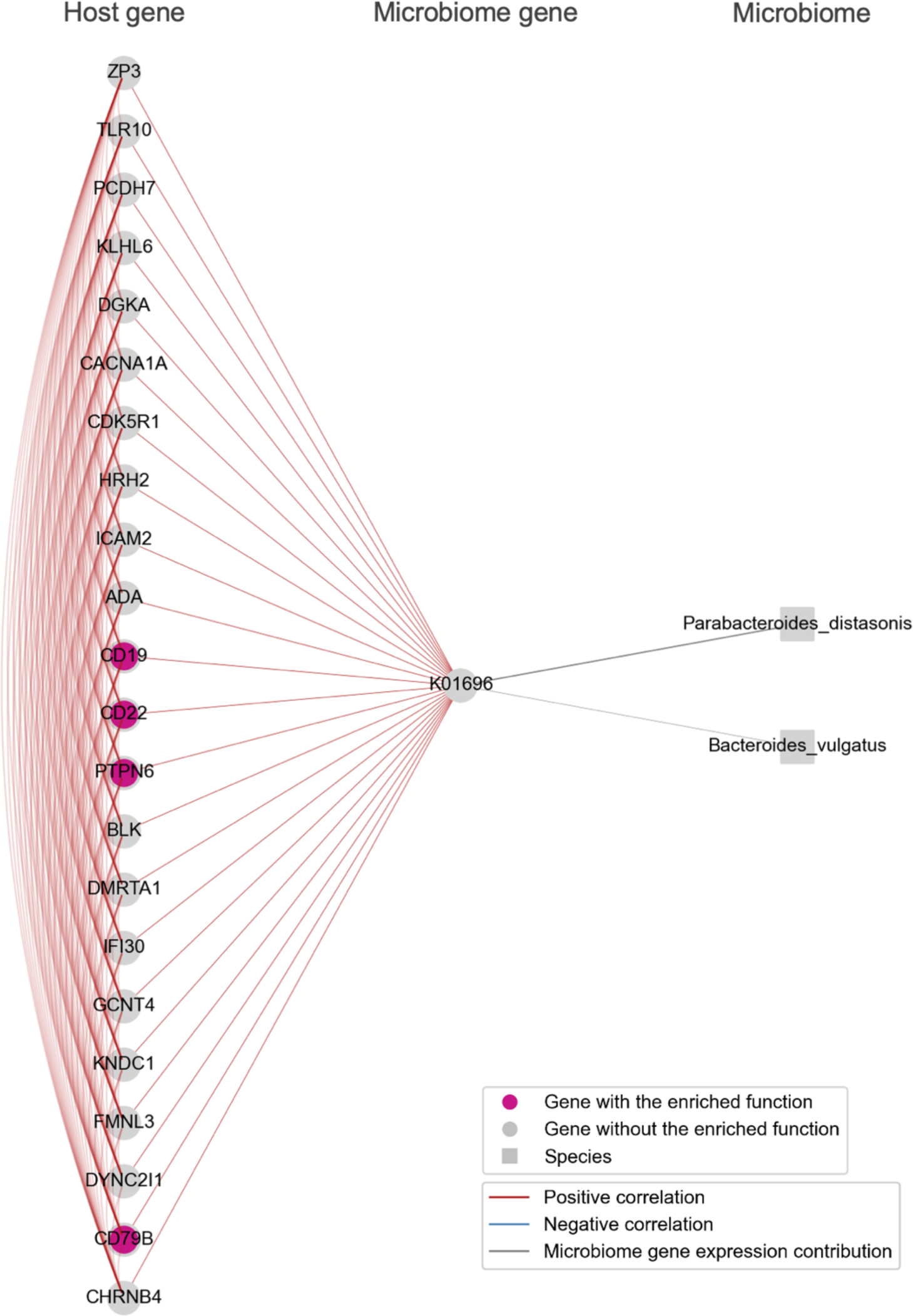
B cell receptor signaling pathway-enriched Module_01. Host genes involved in the immune response of B cells, including *CD19*, *CD22*, *CD79B* and *PTPN6*, which were highly expressed in the cecum, co-expressed with the K01696 (*trpB*). These genes were visualized alongside bacterial species nodes that contribute to the expression of the microbiome genes. K01696 (*trpB*) expression was contributed to by *Parabacteroides distasonis*. Edges among genes indicate an absolute Spearman’s correlation coefficient greater than 0.8. Edges connecting microbiome genes to bacterial species are shown when the contribution rate of bacterial species to microbiome gene expression exceeds 0.2. Note that the bacterial species nodes were added after the module detection analysis. The thickness of the edge increases with higher contribution rates.

**Fig. 7.**
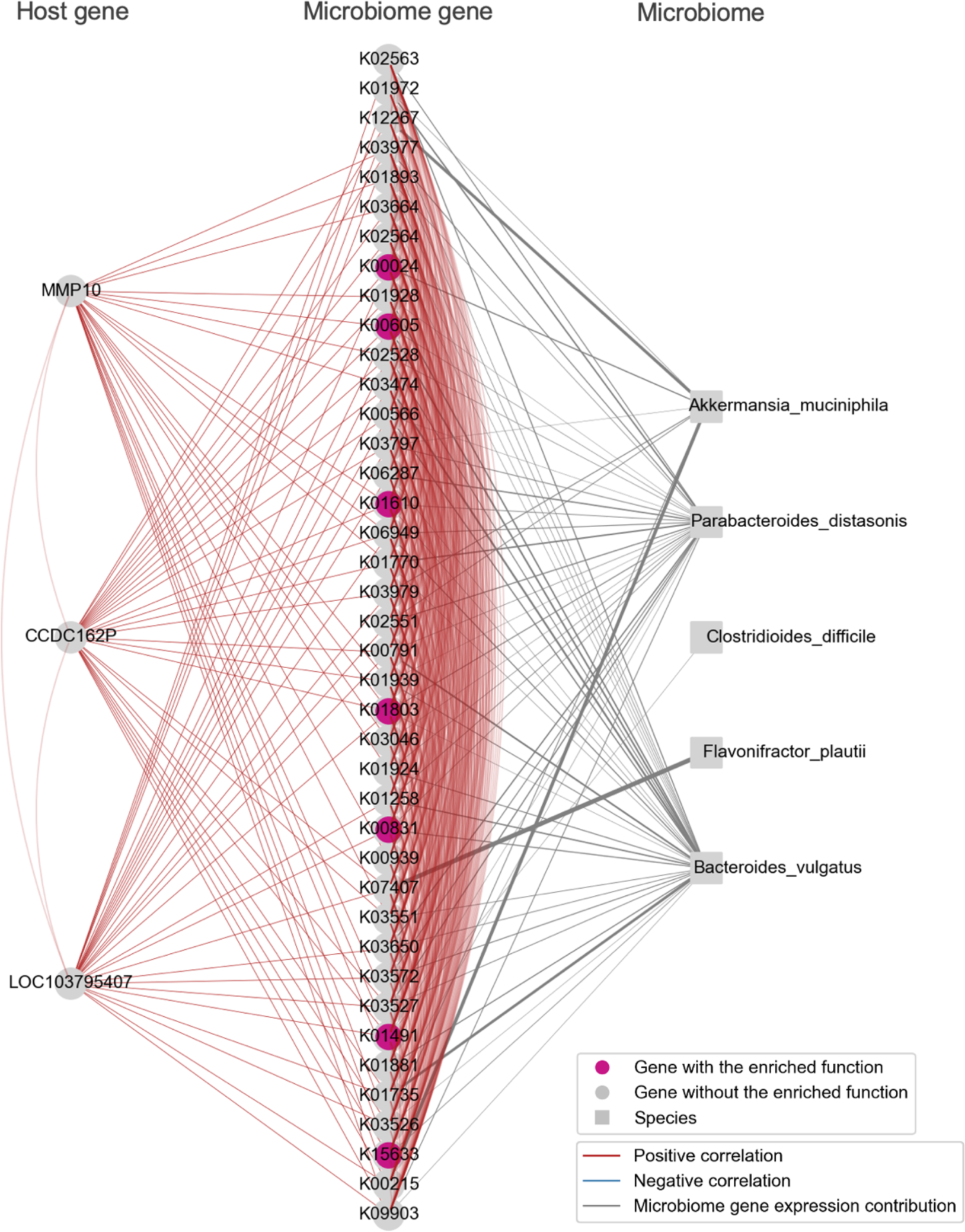
Carbon metabolism pathway-enriched Module_21. Host genes, including *MMP10*, which is specifically highly expressed in the cecum, and 40 microbiome genes involved in carbon metabolism, were identified. These genes were visualized alongside bacterial species nodes that contribute to the expression of the microbiome genes. Note that the bacterial species nodes were added after the module detection analysis. Edges among genes indicate an absolute Spearman’s correlation coefficient greater than 0.8. Edges connecting microbiome genes to bacterial species are shown when the contribution rate of bacterial species to microbiome gene expression exceeds 0.2. The thickness of the edge increases with higher contribution rates.

**Fig. 8.**
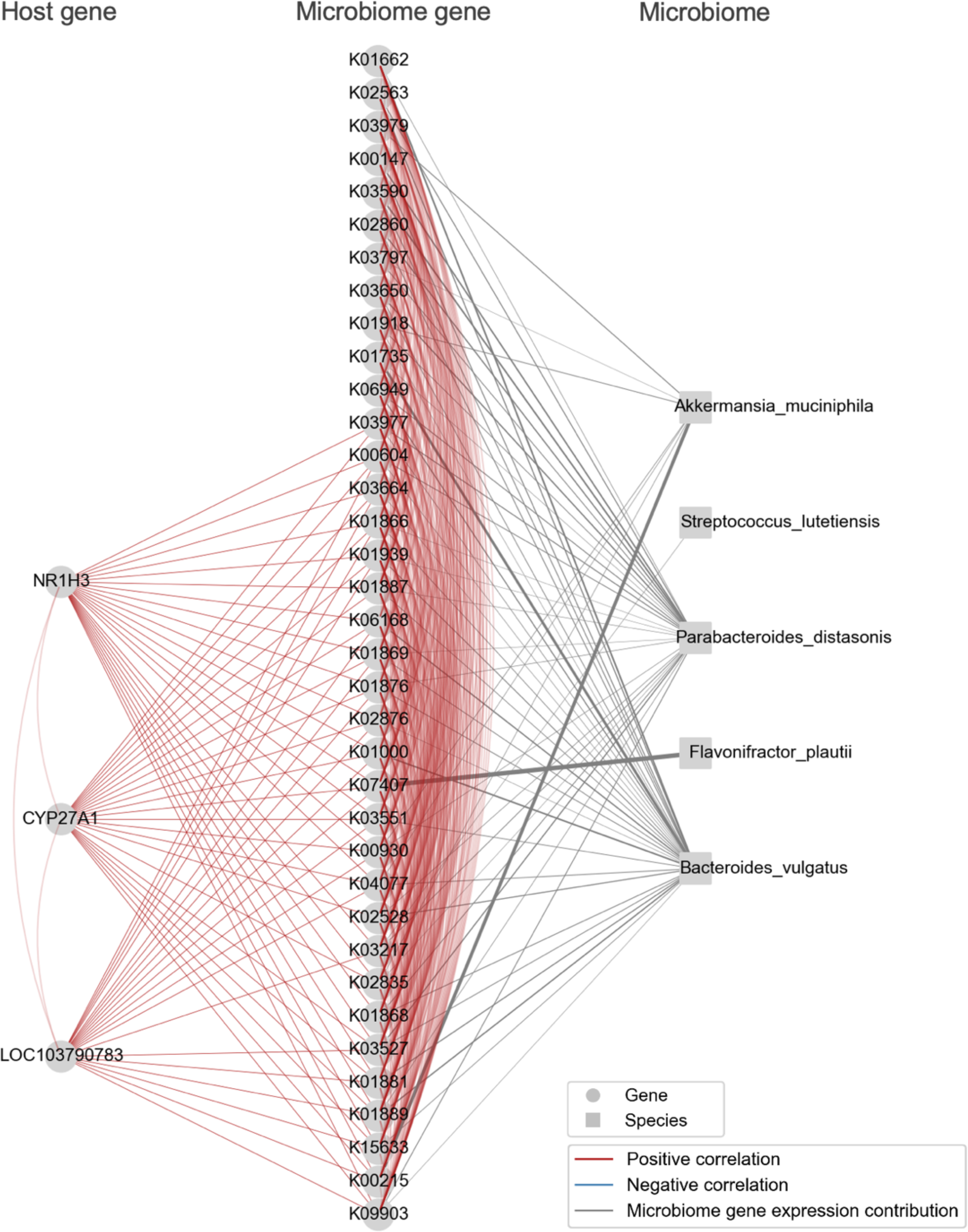
Module_18 containing host genes and microbiome genes related to the production and breakdown of bile acids. These genes were visualized alongside bacterial species nodes that contribute to the expression of the microbiome genes. Note that the bacterial species nodes were added after the module detection analysis. Edges among genes indicate an absolute Spearman’s correlation coefficient greater than 0.8. Edges connecting microbiome genes to bacterial species are shown when the contribution rate of bacterial species to microbiome gene expression exceeds 0.2. The thickness of the edge increases with higher contribution rates.

Module_07, as shown in Figure 5, was identified as a module with a positive correlation between the host gene Fructose-1,6-bisphosphatase 1 (*FBP1*), which is highly expressed in the transverse colon, and 40 microbiome genes. These 40 gut microbiome genes included those associated with fructose-mannose metabolism, such as K01840 (*manB*), K00850 (*pfkA*), and K01818 (*fucI*), as well as those related to oxidative phosphorylation, such as K00241 (*sdhC*, *frdC*), K00426 (*cydB*), K02112 (*atpD*), and K02121 (*ntpE*, *atpE*). The most substantial contributor to the expression of these microbiome genes was *Bacteroides vulgatus*.

*FBP1* is known to have reduced expression in various tumor tissues, including the esophagus (30), liver (31), lung (32), and ovarian cancer (33). It has been confirmed that its expression increases with fructose treatment in intestinal organoids (34). Previous studies have identified *FBP1* as a differentially expressed gene in the small intestine of germ-free mice, suggesting that the intestinal microbiome influences the expression of the host’s *FBP1* gene (35), but the bacterial species and genes affecting the expression of the *FBP1* gene have not been identified. *Bacteroides vulgatus*, which contributed to the expression of fructose-mannose metabolism-related microbiome genes observed in Module_07, has been reported to release fructose by decomposing polysaccharides (36, 37). In this module, the gene K00850 (*pfkA*) involved in the fructo-oligosaccharide degradation and the gene K01818 (*fucI*) involved in the 2’-fucosyllactose degradation were detected, revealing a novel association between polysaccharide degradation genes encoded by *Bacteroides vulgatus* and the host gene *FBP1*.

In Module_01, the B cell receptor signaling pathway was enriched (Fig. 6). This module featured covariation between host B-cell specific genes, including *CD19*, *CD22*, *CD79B*, and *PTPN6*, which were highly expressed in the cecum, and the tryptophan synthesis gene K01696 (*trpB*) from the microbiome. The upregulation of these host B-cell specific genes indicates an increase in B cells within the cecum.

This tryptophan synthesis gene K01696 (*trpB*) expression was contributed to by *Parabacteroides distasonis*. Tryptophan is an essential amino acid that, upon uptake by intestinal cells and subsequent metabolism, can activate the Aryl hydrocarbon receptor and influence B cell immune responses (38). Tryptophan is also known to ameliorate autoimmune diseases, such as inflammatory bowel disease (39). *Parabacteroides distasonis* has been suggested to be involved in various autoimmune conditions in several studies (40, 41). For instance, *Parabacteroides distasonis* levels have been shown to be lower in multiple sclerosis (MS) patients compared to healthy controls, and transplanting the intestinal microbiota from MS patients to germ-free mice resulted in increased severity of autoimmune encephalomyelitis symptoms (40). While previous studies have reported associations between tryptophan and autoimmune diseases via B-cell immune responses, as well as between autoimmune diseases and *Parabacteroides distasonis*, this study identifies a correlation between the tryptophan synthesis gene K01696 (*trpB*) from *Parabacteroides distasonis* and B-cell specific genes (*CD19*, *CD22*, *CD79B*, and *PTPN6*).

In Module_21, a group consisting of three host genes and 40 microbiome genes was identified. The three host genes included *MMP10*, two uncharacterized genes LOC103795407 and *CCDC162P*, all of which showed high expression in the cecum (Fig. 7). The microbiome genes that showed a positive correlation with these genes included K01491 (*folD*), K00605 (*gcvT*), K01803 (*tpiA*), K00831 (*serC*), K15633 (*gpmI*), K00024 (*mdh*), all of which are involved in carbon metabolism pathways. *Bacteroides vulgatus* was identified as the bacteria contributing to the expression of these microbiome genes. The *MMP10* gene plays a crucial role in maintaining epithelial barrier function and is necessary for resolving colonic epithelial damage (42). A study examining the host transcriptome and microbiota composition of colorectal cancer tumors found high expression of *MMP10* in tumor tissues and a significant increase in the genus *Bacteroides*, indicating a link between the host gene *MMP10* and the genus *Bacteroides* (43). These findings are consistent with our observations. Furthermore, in the identified Module_21, not only the relationship between the host gene *MMP10* and *Bacteroides vulgatus*, but also a group of microbiome genes related to carbon metabolism pathways showing a positive correlation with the host gene *MMP10* was identified. One hypothesis explaining the observed positive correlation between carbon metabolism-related genes encoded by *Bacteroides vulgatus* and the host gene *MMP10* is that the microbiome may form colonies within the host’s mucosal layer. Mucin, the principal component of mucus secreted by the gastrointestinal mucosal epithelium, contains galactose and promotes bacterial settlement and adhesion (44), serving as a nutrient source for certain bacteria. *Bacteroides vulgatus* possesses mucosal adherence factors, allowing access to host-derived carbon sources (45). The upregulation of *MMP10* in the cecum might represent a protective response against colonization by *Bacteroides vulgatus*. The detection of carbon metabolism-related genes of *Bacteroides vulgatus* in this module_21 suggests they might be utilizing carbon sources derived from mucin or dietary glucose.

Specifically, while *Bacteroides vulgatus* cannot degrade human mucin independently, it can acquire carbon sources from mucin in co-culture with *Akkermansia muciniphila* (46), which possesses genes detected in this module. When *Akkermansia muciniphila* converts galactose to glucose, *Bacteroides vulgatus* might then activate these carbon metabolism-related genes to utilize this glucose (Fig. S1). Our findings indicate a relationship between the carbon metabolism-related genes of *Bacteroides vulgatus* and the host gene *MMP10*; however, further experiments are required to ascertain whether the upregulation of *MMP10* serves a protective role against colonization by *Bacteroides vulgatus* and *Akkermansia muciniphila* or if *Bacteroides vulgatus* utilizes mucin as a carbon source.

In Module_18, three host genes and 54 microbiome genes were identified, and the expression of these microbiome genes was further influenced by five bacterial species. The relationships between these host genes and microbiome genes, as well as between the host genes and bacterial species, were consistent with existing knowledge about bile acid synthesis and degradation (Fig. 8). This module included the host gene *CYP27A1*, which is involved in bile acid production (47), and *NR1H3*, a host gene that codes for a bile acid receptor (48). These genes were upregulated in the cecum and transverse colon compared to the rectum. The microbiome genes interacting with these host genes included K03664 (*smpB*), a gene essential for bacterial survival when upregulated in the presence of secondary bile acid deoxycholic acid, K02835 (*prfA*), which positively regulates the expression of a gene coding for bile acid hydrolase, and K03527 (*ispH*, *lytB*), a gene involved in bile acid sensitivity. Additionally, *Bacteroides vulgatus* contributed to the expression of all these microbiome genes, which aligns with previous studies showing an increase in *Bacteroides* spp. correlating with the expression of the host gene *NR1H3* (49).

## DISCUSSION

In this study, we extracted gene modules from a host and microbiome gene co-expression network based on dual transcriptome profiling. The module extraction method proposed in this study is based on clique finding and takes into account the structure of three types of networks: host-host, host-microbiome, and microbiome-microbiome. This approach enables the extraction of modules containing both host genes and microbiome genes. Conventional module extraction methods do not differentiate between host gene nodes and microbiome gene nodes within the network, potentially resulting in modules composed exclusively of either host genes or microbiome genes. Indeed, when conventional module extraction methods such as the Newman algorithm (15), the Louvain algorithm (16), the Leiden algorithm (17), and Weighted Correlation Network Analysis (WGCNA) (18) were applied to our dataset (Supplementary Note Section 9), we observed that these modules included those composed solely of host genes or microbiome genes (Table S5). Notably, about half of the genes in WGCNA modules were found in modules comprising only one species. While the purpose of module extraction is to interpret the interactions among genes within a module, these single-species modules do not allow for an interpretation of the relationship between host genes and microbiome genes. Furthermore, to verify whether functionally similar genes were being extracted, we conducted enrichment analysis on the extracted modules (Supplementary Note Section 8). The results showed that our method had the highest proportion of functionally characterized modules (enriched modules) (Table S6).

The gene modules extracted, which encompassed both host and microbiome genes, frequently exhibited relationships where a small number of genes regulated a larger number of genes between the host and microbiome genes. These observations are thought to reflect the characteristics of the interactions between the host and the microbiome. Several studies have reported the impact of host gene mutations on the intestinal microbiome. For instance, it has been reported that the host gene *VDR*, which codes for the Vitamin D receptor, influences the colonization of *Parabacteroides* (50), and that the host gene *ABO* is associated with *Faecalicatena lactaris* (51). In another study investigating the relationship between the intestinal microbiome and host gene loci through quantitative trait locus analysis, it was suggested that *Akkermansia muciniphila* regulates the expression of *ATF3*, which codes for a transcription factor that plays a crucial role in metabolic and immune regulation, and this bacterium is related to only four genes (52). These studies highlight the fact that a single or a small number of host genes are controlling, or are controlled by, the intestinal microbiome. Conversely, a small number of microbiome genes can affect the host’s metabolic (53), fitness (54), and longevity (55) phenotypes. For example, structural mutations in the gene related to butyrate transport in *Anaerostipes hadrus* are strongly associated with a reduction in the host’s metabolic risk (53), and the intestinal microbiome’s thiamine biosynthesis genes, transferred by horizontal transmission, influence the host’s fitness (54). Considering these previous studies and our results, it is suggested that the interaction between the host and the microbiome involves widespread influence of one’s genes on the activity of the other.

The common marmoset used in this study bears anatomical and pharmacological similarities to humans (19), and the fecal microbiome of the common marmoset is more similar to that of humans compared to other animal models (13). While the common marmoset’s cecum—one of the study’s targeted intestinal sites—does not possess an appendage as pronounced as the human appendix, it is known to share significant similarities with the human appendix in terms of microanatomical structure and immune reactivity (56, 57). The results of this study using common marmosets could therefore hold higher applicability to humans compared to other animal models.

In conclusion, we generated host transcriptome and microbiome metatranscriptome profiles of the common marmoset and performed gene co-expression network analysis. The algorithm we proposed for extracting gene modules in the gene co-expression network enabled the identification of dense, functionally related host-microbiome co-expression modules, which was challenging using the conventional module extraction algorithms. The identified gene modules provided new insights into relationships between host genes and intestinal microbiota and microbial genes bridging interactions between the host and bacteria. Our findings underscore that interactions between the host and microbiome are not uniform throughout the colon but are localized, indicating spatial variation in host-microbiome interactions along the colon. These insights could be crucial in understanding mechanisms behind localized gut diseases, such as colorectal cancer and inflammatory bowel disease, and in developing localized therapies (58, 59, 60, 61).

## MATERIALS AND METHODS

### Animal experiments

Common marmosets were housed at the Central Institute for Experimental Animals (Kawasaki, Japan). Five marmosets were euthanized with intravenous administration of pentobarbital overdose, and the digestive tract was isolated (Table S7, Supplementary Note Section 1). Three gastrointestinal site samples (cecum, transverse colon and rectum), including the mucosal layer, were used for dual RNA-seq. These three sites were targeted at the beginning, middle and end of the colon, which has an abundant microbiome. The animal experiment protocol was approved by the CIEA Institutional Animal Care and Use Committee (approval no. 17031, 18032, and 21084). The study was conducted in accordance with the guidelines of CIEA that comply with the Guidelines for Proper Conduct of Animal Experiments published by the Science Council of Japan. Animal care was conducted in accordance with the Guide for the Care and Use of Laboratory Animals (Institute for Laboratory Animal Resources, 2011).

### RNA extraction and dual RNA sequencing of host and microbiome

RNA libraries were prepared after removing rRNA from the extracted high-quality Total RNA (RNA integrity number (RIN) scores ≥8.6; Table S1). Illumina NovaSeq 6000 v1.5 sequencing yielded a total of 472 Gnt of paired-end reads (100 nt × 2). This dataset included an average of 314.9 M reads per sample (Table S2 and S8; Supplementary Note Section 2).

### Construction of host-microbiome co-expression network and identification of gene modules

In this study, we utilized gene co-expression network analysis to identify interacting gene modules of host and microbiome (Supplementary Note Section 7). We established three types of networks: host-host, host-microbiome, and microbiome-microbiome, wherein genes were considered as nodes and co-expressing genes were interconnected (Fig. 1D). Networks were defined by an adjacency matrix, where the adjacency *a_ij_* between gene *i* and gene *j* was defined as follows based on co-expression similarity *s_ij_*:

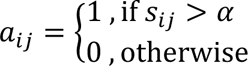

The co-expression similarity *s_ij_* was computed using the Spearman’s correlation coefficient calculated for genes *i* and *j* represented by vectors of TPM values for host-host and microbiome-microbiome networks, and absolute values of Spearman’s correlation coefficients for host-microbiome networks. The adjacency threshold α was set at 0.80. These three types of networks were then integrated into one.

We developed an algorithm to identify densely interacting gene modules from the host-microbiome co-expression network. This algorithm, based on clique finding, progressively merges cliques (Fig. 1E). In graph theory, a “clique” refers to a subgraph within the network where all nodes are fully connected. In the context of a gene co-expression network, all genes within a clique indicate a strong co-expression (correlation) amongst them. In this study, we focus only on networks and cliques that include gene nodes from both the host and microbiome, encompassing three types of networks: host-host, host-microbiome, and microbiome-microbiome. Our algorithm identified modules of interacting genes in the gene co-expression network as aggregates of overlapping cliques. It was implemented in three steps: (1) Cliques within the network were found; (2) Beginning with the largest cliques, we examined their overlap with any other cliques. If overlap occurred, we computed the cliqueness scores independently for each of the three types of networks (host-host, host-microbiome, and microbiome-microbiome networks) upon merging the overlapping cliques. The cliqueness scores for the host-host, host-microbiome, and microbiome-microbiome networks were denoted as *C_hh_*, *C_hm_*, and *C_mm_*, respectively; (3) Cliques were merged if they satisfied the condition: *C_hh_* > τ, *C_mm_* > τ, and *C_mm_* > τ. The threshold τ for cliqueness was set at 0.60 based on parameter determination (Supplementary Note Section 7). If there were multiple cliques with minimum cliqueness scores exceeding the threshold, the clique with the highest cliqueness score was selected. Steps (2) and (3) were repeated until no cliques could be merged.

For the host-host networks the cliqueness score *C_hh_* between cliques A and B was defined as:

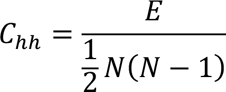

Here, *N* is defined as the number of host gene nodes in the union of cliques A and B, represented as *N* = |*N_A_* ∪ *N_B_*|. *N_A_* and *N_B_* denote the number of host gene nodes within clique A and clique B, respectively. *E* corresponds to the total number of edges between the host nodes included in cliques A and/or B, formulated as *E* = *E_A_* + *E_A_* + *E_AB_*. Here, *E_A_* and *E_B_* represent the number of edges between host gene nodes within clique A and clique B, respectively, while *E_AB_* denotes the number of edges connecting across host gene nodes within clique A and clique B. The cliqueness score for microbiome-microbiome networks *C_mm_* was calculated in the same way.

For the host-microbiome network, the cliqueness score *C_hm_*, was defined as:

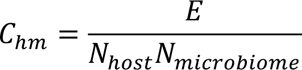

where *N_host_* and *N_microbiome_* represent the number of host and microbiome gene nodes respectively in the union of cliques A and B. *E* corresponds to the total number of edges between the host or microbiome gene nodes included in cliques A and/or B.

## Data and source code availability

All raw sequence data have been submitted to the DDBJ under project PRJDB16352 from the Ministry of Education, Culture, Sports, Science and Technology of Japan. The gene expression profiles of the host and microbiome are available at https://doi.org/10.5281/zenodo.8264866. The algorithm for module extraction is available at https://github.com/MikaUhr/MEIGN.git.

## ACKNOWLEDGEMENTS

We thank the scientists of the Central Institute for Experimental Animals for providing support for animal experiments and the members of the Sakakibara Lab at Keio University for helpful discussions.

This work was supported by grants from the Japan Agency for Medical Research and Development (AMED PRIME) Grant Number JP19gm6010006 and JST, CREST Grant Number JPMJCR20S3, Japan. M. Uehara has received funding from JSPS KAKENHI Grant Numbers JP20J21477.

M.U. performed the experiments, conducted the bioinformatics analysis and cowrote the paper; T.I. and E.S. provided common marmoset samples; M.U. and S.H. performed RNA extraction; A.T. performed deep sequencing with high-throughput sequencer; Y.S. designed and supervised the research, analysed the data, and cowrote the paper. All authors have read and approved the manuscript.

The authors declare no competing interests.

## Supplemental Materials

**Supplementary Notes.** Supplementary information on the methods, consisting of: Section 1. Animal experiments; Section 2. RNA extraction and dual RNA sequencing of host and microbiome; Section 3. Partitioning of host RNA reads and microbiome RNA reads; Section 4. Accuracy assessment of partitioning of RNA read data into host and microbiome reads; Section 5. Analysis of host gene expression; Section 6. Analysis of microbiome gene expression; Section 7. Network construction and determination of parameters for module extraction; Section 8. Characterization and evaluation of identified modules; Section 9. Comparison with other module extraction methods

**Table S1.** RIN score of total RNA samples

**Table S2.** Sequence statistics

**Table S3.** Number of RNA reads derived from the host and microbiome

**Table S4.** Results of DEG analysis with pairwise DEG analysis between three intestinal sites

**Table S5.** Statistics on the module extraction methods

**Table S6.** Evaluation of the module extraction methods through enrichment analysis

**Table S7.** Information on common marmoset samples

**Table S8.** Number of RNA reads at each preprocessing step

**Fig. S1.** Carbon metabolism-related microbiome genes detected in Module_21

The Carbon metabolism pathway of KEGG is shown with the genes in Module_21 highlighted in red.

**Fig. S2.** Changes in the coefficient of determination for fitting to the scale-free topology model in response to the soft threshold in WGCNA

The red line indicates a coefficient of determination = 0.8.

