## Supplementary Note; Supplementary Table S1, S2, S3, S4, S5, S6, S7 and S8; Fig. S1, S2 for "Decoding host-microbiome interactions through co-expression network analysis within the non-human primate intestine"

### Supplementary Notes

#### Section 1. Animal experiments

Common marmosets were housed at the Central Institute for Experimental Animals (Kawasaki, Japan) with free access to a pellet diet (New World Monkey Diet, CMS-1M; CLEA Japan, Tokyo, Japan). The cage size was 820 × 610 ×1578 mm, and the cages were positioned facing each other to allow the animals to communicate visually and vocally. All cages were equipped with a sleeping area, wooden perches, and hammocks. The animal rooms were maintained at 26–28 °C and 40–60% humidity with a 12 h:12 h light/dark cycle. The animals were negative for Salmonella spp., Shigella spp. and Yersinia pseudotuberculosis in yearly fecal examinations. Five marmosets were euthanized with intravenous administration of pentobarbital overdose, and the digestive tract was isolated (Table S7). The gastrointestinal tract of each animal was excised, and each intestinal site was collected. The samples of the intestinal site were immediately frozen in liquid nitrogen and stored at -80 °C. The samples were crushed and homogenized in solution D containing guanidinium, which inhibits ribonuclease (1), within one week after dissection to protect against degradation and stored at -80 °C. Three gastrointestinal site samples (cecum, transverse colon and rectum), including the mucosal layer, were used for dual RNA-seq. These three sites were targeted at the beginning, middle and end of the colon, which has an abundant microbiome.

The animal experiment protocol was approved by the CIEA Institutional Animal Care and Use Committee (approval no. 17031, 18032, and 21084). The study was conducted in accordance with the guidelines of CIEA that comply with the Guidelines for Proper Conduct of Animal Experiments published by the Science Council of Japan. Animal care was conducted in accordance with the Guide for the Care and Use of Laboratory Animals (Institute for Laboratory Animal Resources, 2011).

#### Section 2. RNA extraction and dual RNA sequencing of host and microbiome

RNA was extracted by a combination of the acid-guanidium-phenol-chloroform RNA extraction method (2) and bead crushing method and assessed to ensure high quality (RNA integrity number (RIN) scores ≥8.6; Table S1). rRNA was removed using the Ribo-Zero Plus rRNA Depletion Kit (Epidemiology) (Illumina). Sequencing libraries were prepared using the TruSeq Stranded Total RNA HT Kit (Illumina). All these procedures were performed according to the manufacturer's instructions. Illumina NovaSeq 6000 v1.5 sequencing yielded a total of 472 Gnt of paired-end reads (100 nt × 2). This dataset included an average of 314.9 M reads per sample (Table S2). The libraries were adapter trimmed and quality filtered using fastp version 0.20.1 (3) with "--length_required 50, --low_complexity_filter, --correction, --trim_poly_x --qualified_quality_phred 20 --average_qual 27". The rRNA reads were removed by SortMeRNA (4) version 2.1 with "-e 1e-10" (Table S8).

#### Section 3. Partitioning of host RNA reads and microbiome RNA reads

Dual RNA-seq data, which contains a mixture of RNA reads originating from the host and the microbiome, necessitates the separation of host and microbiome RNA. We mapped the paired-end reads to the common marmoset reference genome (Callithrix jacchus; GenBank assembly accession GCA_009663435.2) using HISAT2 version 2.1.0 (5) with "--mp 3, 2, --score-min -0.7". Reads that mapped to the host genome were considered host derived (Table S3). Unmapped reads were then subjected to single-end mapping with the same parameters to prevent potential contamination of marmoset-derived reads in the microbiome read dataset. Only those reads for which neither forward read 1 nor read 2 mapped to the host genome were considered microbiome derived. The method for evaluating the accuracy of this partitioning of host-derived and microbiome-derived RNA reads is described in the Supplementary Note Section 4.

**Section 4. Accuracy assessment of partitioning of RNA read data into host and microbiome reads**

The pipeline developed in this study separates host-derived reads from microbial community-derived reads in dual RNA-seq data by aligning the RNA reads to the genome of the host, the common marmoset. We therefore prepared simulation datasets and conducted accuracy evaluations for separating host-derived reads and microbial community-derived reads. The simulation data were based on the Coding DNA Sequence (CDS) of the common marmoset and the microbial community. For the microbial community CDS, the top 32 species were chosen based on their relative abundance across all samples. These 32 species cover 70.2% of the total (Table S16 posted at https://doi.org/10.5281/zenodo.8264876). Using these CDS sequences, we generated 100 bp paired-end reads for the host and the microbial community, producing 10M reads and 1M reads randomly, and created five datasets using the tool wgsim version 0.3.1 (https://github.com/lh3/wgsim). The error rate in the simulation data followed that of NovaSeq 6000. Using these datasets, we employed the splice-aware aligner HISAT2 and adjusted the mismatch penalty scores with --mp and the alignment threshold with --score-min. The parameter combination that resulted in the highest F1 score was "--mp 3,2, --score-min -0.7". We defined the number of marmoset reads classified as marmoset reads as true positives (TP), the number of bacterial reads classified as marmoset reads as false positives (FP), and the number of marmoset reads not classified as marmoset reads as false negatives (FN). We then calculated precision (TP/(TP+FP)), recall (TP/(TP+FN)), and F-score (F=2 × ((precision × recall)/(precision +recall))) (Table S9 posted at https://doi.org/10.5281/zenodo.8264876).

**Section 5. Analysis of host gene expression**

The BAM files of RNA reads mapped to the common marmoset genome were sorted using SAMtools version 1.3.1 (6). Gene-level read counts were then calculated using HTSeq version 0.11.2 (7). Gene expression profiles were normalized to transcripts per million (TPM). We used 16,578 genes, whose expression was confirmed in all 15 samples across three sites from five individuals, for downstream analysis. Differentially expressed genes (DEGs) between intestinal sites were analyzed using the R package DESeq2 version 1.40.1 (8). We conducted pairwise Wald tests among the three sites and adjusted the resulting P-values using the Benjamini-Hochberg method for multiple testing corrections. Genes with an FDR $< 0.01$ were identified as DEGs. Further, we performed enrichment analysis for all host genes across three sites in pairwise comparisons using the KEGG database (release 94.1) (9) and GSEA version 4.3.2 (10). This approach enabled the identification of site-specific KEGG pathways (normalized enrichment score $>1.5$, FDR$<0.1$). While GSEA typically uses a threshold of FDR $< 0.25$, we opted for a threshold of FDR $< 0.1$ to further reduce the potential for false positives (https://www.gsea-msigdb.org/gsea/doc/GSEAUserGuideFrame.html?Run_GSEA_Page).

**Section 6. Analysis of microbiome gene expression**

Functional profiling of microbiome RNA reads was performed using HUMAnN3 version 3.0.1 (11), which identifies species profiles, aligns RNA reads to the pangenome, and conducts translation searches on unclassified reads to quantify gene expression levels. Samples with more than 500,000 reads annotated with KEGG Orthology (KO) functions (9) were used in downstream analyses (Table S17).

**Section 7. Network construction and determination of parameters for module extraction**

The gene co-expression network was constructed from host genes identified as DEGs among the three intestinal sites and microbiome genes expressed (copies per million (CPM) > 0) in 10 or more samples. Due to the nature of dual RNA-seq, the microbiome gene expression profiles tend to be sparser with fewer reads compared to the host. Genes with a high prevalence of zero expression values could introduce false correlations. However, there is also a possibility that genes showing low expression might be incorrectly recorded as having zero expression. Therefore, to retain genes that might have low expression in one of the three sites but are erroneously recorded as having zero expression, we included microbiome genes confirmed to be expressed in more than ten samples.

Our method requires the selection of gene co-expression network types and the cliqueness score threshold for merging cliques. As described in the subsection "Construction of host-microbiome co-expression network and identification of gene modules" in the MATERIALS AND METHODS section, the gene co-expression networks were defined using adjacency matrices. We constructed two network types: "the positive network" and "the mixed network". The positive network, as mentioned in the text, used Spearman's correlation coefficients for host-host and microbiome-microbiome networks and the absolute value of Spearman's correlation coefficients for host-microbiome networks. On the other hand, the mixed network was calculated using the absolute values of Spearman's correlation coefficients for all three networks. While the Positive network draws edges between gene nodes with positive correlations in the host-host and microbiome-microbiome networks, the mixed network draws edges between gene nodes with either positive or negative correlations. The choice of Spearman's correlation coefficient over Pearson's correlation coefficient was due to the possibility of our data not following a normal distribution. The clique-based approach is a method for extracting modules by detecting cliques, which are subgraphs where all nodes are fully connected. To achieve this, it is necessary to convert the problem into a binary question of whether there is an edge between nodes (i.e., whether they are connected) or not, as opposed to considering a weighted graph. Therefore, we employed the threshold of 0.8 for Spearman's correlation coefficient. Generally, a correlation value exceeding 0.8 is described as a strong correlation. In the context of gene co-expression, this threshold has been demonstrated to be suitable for predicting genes with similar functions and identifying genetic interactions (12, 13).

The threshold for the cliqueness score was used to determine the merging of overlapping cliques in the host-host, host-microbiome, and microbiome-microbiome networks. These score thresholds were set in increments of 0.1, ranging from 0.3 to 0.9. The extracted gene modules were subjected to KEGG pathway enrichment analysis as described in the Supplementary Note Section 8. The highest proportion of enriched communities was observed when applying a cliqueness score threshold of 0.60 to the positive network. Based on these results, module extraction was further performed with cliqueness scores ranging from 0.51 to 0.69 in increments of 0.01. Ultimately, a threshold of 0.60 was determined to be optimal (Table S18 posted at https://doi.org/10.5281/zenodo.8264876).

**Section 8. Characterization and evaluation of identified modules**

The modules ultimately obtained were subjected to an enrichment analysis using Fisher's exact test with Benjamini-Hochberg multiple testing correction for the interpretation of gene functions within the modules and evaluation of the module extraction methodology. The modules were functionally characterized by KEGG pathways (FDR<0.05; Table S15), both for interpretation of gene functions within modules and for evaluation of module extraction methodology.
The modules subjected to enrichment analysis were defined as those containing both host and microbiome genes and having a node count ranging from 5 to 500 (hereafter referred to as "target modules"). Module extraction was performed with the objective of interpreting the interactions among genes within a module; therefore, it was necessary to control the number of genes included in each module. Consequently, we set a threshold for the number of genes in target modules to be greater than 10. The enrichment analysis was conducted on all genes within these modules, without differentiating between host and microbiome genes.
We utilized this module-based enrichment analysis for parameter evaluation. This analysis was based on the assumption that functionally similar genes would have co-variant expression levels (14). We selected the parameter that maximized the proportion of enriched modules within the target modules, a metric we have termed as the "rate of enriched modules" (Table S18 posted at https://doi.org/10.5281/zenodo.8264876). This metric is commonly utilized to assess module extraction methodologies and was employed in this study for the comparison of parameters and techniques. In addition to this metric, we also calculated two further indices: "rate of genes in enriched modules" and "rate of enriched genes". The rate of genes in enriched modules indicates the proportion of genes within the enriched modules relative to the total number of genes in all target modules. On the other hand, the rate of enriched genes denotes the proportion of genes that are not only part of the enriched modules but also possess enriched functions, again in relation to the total gene count in all target modules.

**Section 9. Comparison with other module extraction methods**

In this study, the module extraction method we proposed was compared with Newman algorithm (15), Louvain algorithm (16), Leiden algorithm (17), and weighted correlation network analysis (WGCNA) (18), as shown in Table S5 and S6. Newman, Louvain, and Leiden algorithms are designed to extract modules from constructed networks. The Newman and Louvain algorithms adopt an approach that maximizes modularity. The Leiden algorithm, an improvement over the Louvain algorithm, also handles a quality function known as the Constant Potts Model (CPM) (19) in addition to maximizing modularity. WGCNA encompasses a series of processes from network construction to module extraction, defining modules by transforming the similarity matrix of a weighted network into a distance matrix and implementing hierarchical clustering.
For the networks used in Newman, Louvain, and Leiden algorithms, we utilized two types: positive networks and mixed networks as described in the Supplementary Note Section 7. Unlike our method, these algorithms can handle weights, so we tested both weighted and unweighted edges for each network type. A threshold of 0.8 was used for Spearman's correlation coefficient. We experimented with various combinations of network types, edge weights, and algorithm parameters. As detailed in the Supplementary Note Section 8, we used the proportion of enriched modules (FDR < 0.05) obtained from enrichment analysis as an evaluation metric. Ultimately, we selected the combination of parameters that showed the highest evaluation metric (Table S19, S20, and S21). The specifics for each module's parameters are as follows:

- Newman algorithm was implemented using the greedy_modularity_communities module in NetworkX version 3.1. The resolution parameter, determining module size, was tested from

1.0 to 5.0 in 1.0 increments. For the weighted Positive network, the highest evaluation metrics were observed at resolutions of 3.0 and 4.0. Consequently, the resolution was further refined (from 2.1 to 4.9 in 0.1 increments), with 3.4 being the final resolution chosen.

- Louvain algorithm was implemented using the louvain_communities module in NetworkX version 3.1. The resolution parameter was tested in the same manner as Newman's algorithm. Five seed values (111, 112, 113, 114, 115) were used. For the unweighted Positive network, the highest evaluation metric was observed with seed 112 and resolution 3.0. The resolution was further refined (from 2.1 to 3.9 in 0.1 increments), with 3.1 ultimately being selected.
- Leiden algorithm was implemented using Leidenalg version 0.7.0. Two partition types were used: ModularityVertexPartition for maximizing modularity and CPMVertexPartition for utilizing the CPM function. The resolution parameter was tested from 1.0 to 9.0 in 1.0 increments, and five seed values were used. For the unweighted Positive network, the highest evaluation metric was achieved using the CPM function, seed 115, and resolution 8.0. The resolution was further refined (from 7.1 to 8.9 in 0.1 increments), with 8.0 being the final choice.

For WGCNA module extraction, since network construction is part of the method, we followed the WGCNA protocol. WGCNA version 1.72.1 was used. Specifically, when $a_{ij}$ represents the absolute value of the Spearman's correlation coefficient of gene expression between gene $i$ and $j$, the similarity $s_{ij}$was calculated by $s_{ij} = a_{ij}^{\beta}$. Here, $\beta$ represents the soft threshold. Networks were constructed with soft threshold $\beta$ ranging from 1 to 32 in increments of 1, checking for fit to the scale-free topology model. The plot of soft threshold $\beta$ against the coefficient of determination is shown in Fig. S2. The soft threshold $\beta$ was determined to be 21 when the coefficient of determination exceeded 0.8. Subsequently, as the coefficient of determination declined at a soft threshold $\beta$ of 32, networks were constructed with soft thresholds $\beta$ ranging from 21 to 31 in increments of 1. The topological overlap measure (TOM) was calculated from these weights. The distance $d_{TOM_{ij}}$ between genes i and j was calculated as $d_{TOM_{ij}} = 1 - TOM_{ij}$, and modules were extracted through hierarchical clustering. Clustering was performed with all possible values of DeepSplit (ranging from 0 to 4) to control the granularity of cluster division. Finally, the combination of soft threshold and DeepSplit that yielded the highest proportion of enriched modules was selected (Table S22).

These methods, when extracting modules, do not distinguish between host gene nodes and microbiome gene nodes, which can result in modules comprised of only a single species. In this study, based on the assumption that functionally similar genes are co-expressed (20), modules were extracted from the gene co-expression network with the aim of interpreting interactions between host genes and microbiome genes within these modules. Therefore, modules composed solely of a single species were excluded and not included in the evaluation of the method.

**Supplementary Table**

Table S1. RIN score of total RNA samples

| Site | Individual | RIN |
|---|---|---|
| Cecum | CJ01 | 8.6 |
| | CJ02 | 9.5 |
| | CJ03 | 9.7 |
| | CJ04 | 9.9 |
| | CJ05 | 9.9 |
| Transverse colon | CJ01 | 8.8 |
| | CJ02 | 9.2 |
| | CJ03 | 9.8 |
| | CJ04 | 9.9 |
| | CJ05 | 9.7 |
| Rectum | CJ01 | 8.8 |
| | CJ02 | 9.8 |
| | CJ03 | 9.9 |
| | CJ04 | 10.0 |
| | CJ05 | 9.9 |

Table S2. Sequence statistics

| Site | Individual | Number of read pairs | Yield /Gb |
|---|---|---|---|
| Cecum | CJ01 | 82,806,614 | 16.6 |
| | CJ02 | 83,082,185 | 16.6 |
| | CJ03 | 83,900,454 | 16.8 |
| | CJ04 | 79,596,993 | 15.9 |
| | CJ05 | 77,732,809 | 15.5 |
| Transverse colon | CJ01 | 80,584,186 | 16.1 |
| | CJ02 | 104,457,990 | 20.9 |
| | CJ03 | 87,493,619 | 17.5 |
| | CJ04 | 92,702,309 | 18.5 |
| | CJ05 | 126,919,012 | 25.4 |
| Rectum | CJ01 | 302,711,148 | 60.5 |
| | CJ02 | 297,544,429 | 59.5 |
| | CJ03 | 293,969,210 | 58.8 |
| | CJ04 | 235,603,170 | 47.1 |
| | CJ05 | 332,646,363 | 66.5 |

Table S3. Number of RNA reads derived from the host and microbiome

| Site | Individual | Host | Microbiome |
|---|---|---|---|
| Cecum | CJ01 | 54,427,836 | 3,533,996 |
| | CJ02 | 51,135,576 | 4,161,402 |
| | CJ03 | 32,305,866 | 6,643,083 |
| | CJ04 | 55,554,579 | 2,344,913 |
| | CJ05 | 42,038,130 | 3,352,063 |
| Transverse colon | CJ01 | 50,836,124 | 2,650,012 |
| | CJ02 | 70,503,007 | 2,702,415 |
| | CJ03 | 72,043,206 | 414,630 |
| | CJ04 | 67,177,722 | 1,156,958 |
| | CJ05 | 89,558,808 | 1,743,254 |
| Rectum | CJ01 | 225,312,167 | 2,333,324 |
| | CJ02 | 202,417,455 | 2,624,894 |
| | CJ03 | 246,544,903 | 2,348,056 |
| | CJ04 | 187,423,696 | 2,772,388 |
| | CJ05 | 266,283,667 | 2,756,565 |

Table S4. Results of DEG analysis with pairwise DEG analysis between three intestinal sites

| | | Down regulated in | | |
|---|---|---|---|---|
| | | Cecum | T.colon | Rectum |
| Up regulated in | Cecum | - | 307 | 924 |
| | T.colon | 74 | - | 521 |
| | Rectum | 1080 | 776 | - |

Table S5. Statistics on the module extraction methods

| Algorithm | Number of target modules | Number of enriched modules | Number of modules consisting only of a single species | Number of genes in modules consisting only of a single species |
|---|---|---|---|---|
| Proposed method | 27 | 19 | - | - |
| WGCNA | 12 | 8 | 41 | 1632 |
| Leiden | 14 | 8 | 11 | 223 |
| Louvain | 11 | 3 | 10 | 104 |
| Newman | 13 | 6 | 7 | 72 |

Table S6. Evaluation of the module extraction methods through enrichment analysis

| **Algorithm** | **Rate of enriched modules** | **Rate of genes in enriched modules** | **Rate of enriched genes** |
|---|---|---|---|
| Proposed method | **0.704** | 0.761 | **0.281** |
| WGCNA | 0.667 | **0.771** | 0.199 |
| Leiden | 0.571 | 0.665 | 0.012 |
| Louvain | 0.462 | 0.463 | 0.124 |
| Newman | 0.333 | 0.367 | 0.130 |

Table S7. Information on common marmoset samples

| **Notation in this paper** | **Individual ID** | **Sex** | **Date of birth** | **Date of sampling** | **Weight /g** | **Animal experimental approval number** |
|---|---|---|---|---|---|---|
| CJ01 | I6289M | Male | 2015/04/15 | 2017/10/18 | 360 | 17031 |
| CJ02 | I6027M | Male | 2014/02/20 | 2017/12/11 | 306 | 17031 |
| CJ03 | I6421M | Male | 2016/01/08 | 2018/07/04 | 323 | 18032 |
| CJ04 | I7514M | Male | 2020/05/10 | 2021/11/10 | 320 | 21084 |
| CJ05 | I7499M | Male | 2020/04/22 | 2021/12/07 | 314 | 21084 |

Table S8. Number of RNA reads at each preprocessing step

| Site | Individual | Number of raw reads | Number of filtered reads | Number of reads after rRNA removal |
|---|---|---|---|---|
| Cecum | CJ01 | 82,806,614 | 70,283,371 | 59,031,835 |
| | CJ02 | 83,082,185 | 69,882,205 | 55,765,708 |
| | CJ03 | 83,900,454 | 72,486,918 | 39,553,276 |
| | CJ04 | 79,596,993 | 65,889,521 | 58,540,870 |
| | CJ05 | 77,732,809 | 64,121,629 | 46,188,925 |
| Transverse colon | CJ01 | 80,584,186 | 64,616,869 | 54,582,908 |
| | CJ02 | 104,457,990 | 85,259,906 | 74,461,184 |
| | CJ03 | 87,493,619 | 75,345,635 | 73,028,683 |
| | CJ04 | 92,702,309 | 73,436,506 | 69,187,569 |
| | CJ05 | 126,919,012 | 102,158,744 | 93,094,716 |
| Rectum | CJ01 | 302,711,148 | 243,265,414 | 231,339,232 |
| | CJ02 | 297,544,429 | 228,022,976 | 210,077,078 |
| | CJ03 | 293,969,210 | 261,805,007 | 252,411,337 |
| | CJ04 | 235,603,170 | 201,802,337 | 192,220,682 |
| | CJ05 | 332,646,363 | 286,039,812 | 273,899,301 |

### Supplementary Figure

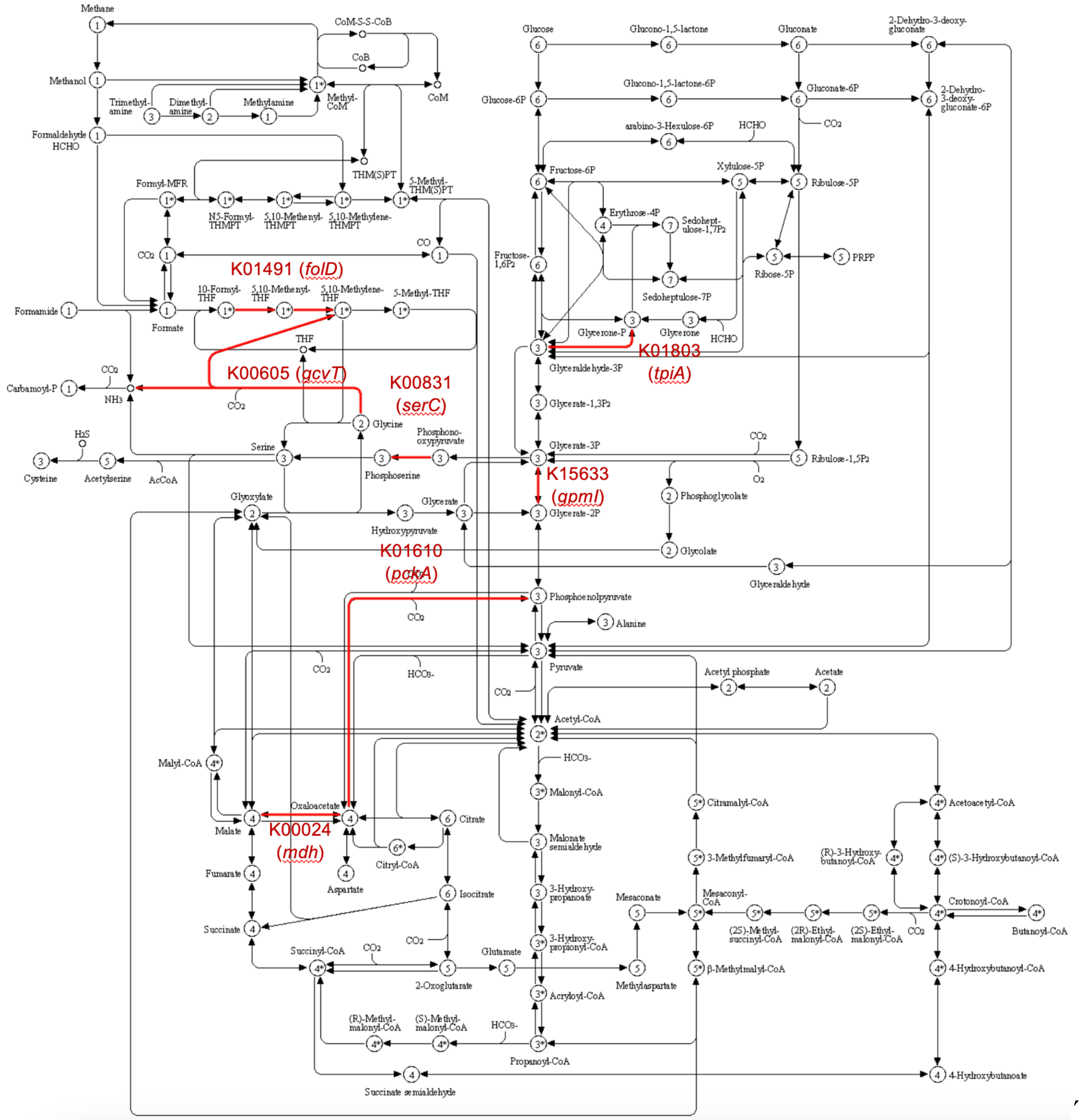


Fig. S1. Carbon metabolism-related microbiome genes detected in Module_21
The Carbon metabolism pathway of KEGG is shown with the genes in Module_21 highlighted in red.

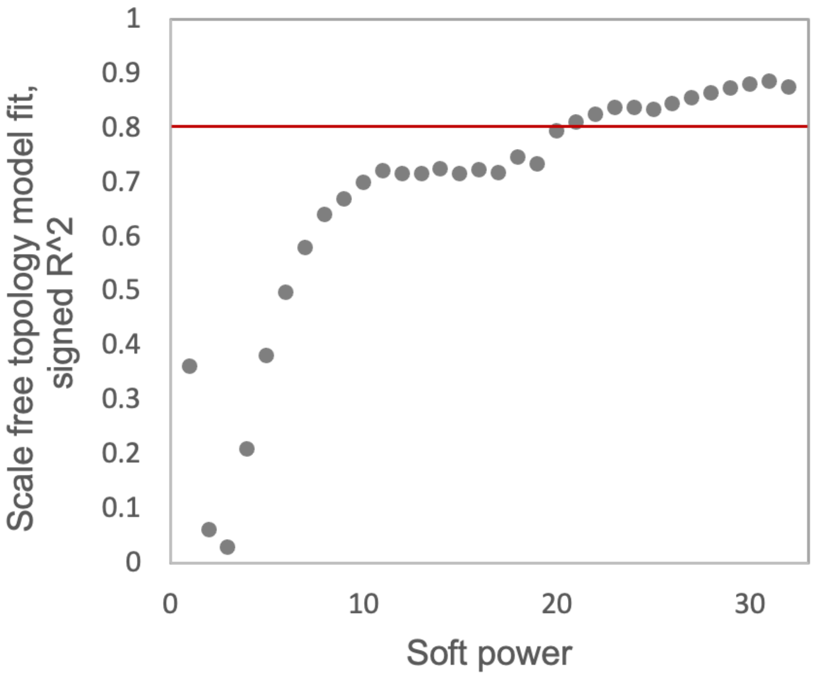


Fig. S2. Changes in the coefficient of determination for fitting to the scale-free topology model in response to the soft threshold in WGCNA

The red line indicates a coefficient of determination = 0.8.
